# Glycomic profiling of the gut microbiota by Glycan-seq

**DOI:** 10.1101/2021.06.30.450488

**Authors:** Lalhaba Oinam, Fumi Minoshima, Hiroaki Tateno

## Abstract

**Background:** There has been immense interest in studying the relationship between the gut microbiota and human health. Bacterial glycans modulate the cross talk between the gut microbiota and its host. However, little is known about these glycans because of the lack of appropriate technology to study them.

**Methods:** We previously developed a sequencing-based glycan profiling method called Glycan-seq, which is based on the use of 39 DNA-barcoded lectins. In this study, we applied this technology to analyze the glycome of the intact gut microbiota of mice. Fecal microbiota was incubated with 39 DNA-barcoded lectins exposed to UV, and the number of released DNA barcodes were counted by next-generation sequencing to obtain a signal for each lectin bound to the microbiota. In parallel, the bacterial composition of the gut microbiota was analyzed by 16S rRNA gene sequencing. Finally, we performed a lectin pull-down experiment followed by 16S rRNA gene sequencing to identify lectin-reactive bacteria.

**Results:** The evaluation of cultured gram-positive (*Deinococcus radiodurans*) and gram-negative (*Escherichia coli*) bacteria showed significantly distinct glycan profiles between these bacteria, which were selected and further analyzed by flow cytometry. The results of flow cytometry agreed well with those obtained by Glycan-seq, indicating that Glycan-seq can be used for bacterial glycomic analysis. We thus applied Glycan-seq to comparatively analyze the glycomes of young and old mice gut microbiotas. The glycomes of the young and old microbiotas had significantly distinct glycan profiles, which reflect the different bacterial compositions of young and old gut microbiotas based on 16S rRNA gene sequencing. Therefore, the difference in the glycomic profiles between young and old microbiotas may be due to their differing bacterial compositions. α2-6Sia-binders bound specifically to the young microbiota. Lectin pull-down followed by 16S rRNA gene sequencing of the young microbiota identified *Lactobacillaceae* as the most abundant bacterial family with glycans reacting with α2-6Sia-binders.

**Conclusion:** The Glycan-seq system can, without any prior culturing and fluorescence labeling, reveal the glycomic profile of the intact bacterial gut microbiota. A combination of lectin pull-down and 16S rRNA gene sequencing can identify lectin-reactive bacteria.

## Background

The microbiota of the digestive tract [1] is dominated by bacteria. It is estimated that 1000 species of commensal, symbiotic, and pathogenic bacteria are present in the gut microbiota [2, 3]. The gut microbiota plays vital roles in human health and disease conditions and is tightly regulated by the lifestyle, dietary habits, and health status of the host [4]. It interacts with the gut epithelium, including the different immune cells within it [5]. The gut microbiota may play regulatory roles in mood, anxiety, and cognition via the gut–brain axis [6], and an imbalance in the gut microbiota may cause gastrointestinal disorders [4, 7] and metabolic [8] and inflammatory diseases [9].

The surface of the bacteria is coated with an intricate network of glycans that act as an interface between mammalian hosts and their gut bacteria [10]. Gram-positive bacterial cells are enclosed by a single membrane covered by a thick peptidoglycan layer and lipoteichoic acids [11], whereas gram-negative bacteria are covered by two cell membranes (inner and outer membranes) separated by a periplasm containing a thin peptidoglycan layer and β-glucan; the outer membrane consists of lipopolysaccharides [12]. Both types of bacteria are often further enclosed by a diverse array of capsular polysaccharides [13]. We previously developed a method to analyze the bacterial cell surface glycomes using a lectin microarray and applied this method to compare 16 different strains of *Lactobacillus casei* [14]. Interestingly, cell surface glycomes differ depending on the bacterial strains. However, there are several drawbacks in the lectin microarray analysis of bacteria, including the following: (1) A large number of cells (0.5 × 10^9^–5 × 10^9^ cells/well) are required for the analysis. (2) Bacteria bound to the lectin microarray are easily released by washing steps meant to remove unbound bacteria. Thus, the results of this analysis may be difficult to reproduce. (3) The analysis requires fluorescently labeled bacterial cells, and different species of bacteria may differ in their fluorescence. As the gut microbiota consists of various bacterial populations, labeling all bacterial populations at the same level of fluorescence is difficult. Hence, the lectin microarray has never been applied to the analysis of the gut microbiota. Thus, despite playing an essential role in bacterial cross talk with the host, bacterial glycans in the gut microbiota remain poorly understood mainly because of insufficient analysis method.

We recently developed a highly multiplexed glycan profiling method called Glycan-seq, which analyzes bulk and single cells using DNA-barcoded lectins and next-generation sequencing [15]. In this study, we applied Glycan-seq to analyze the gut microbiota of mice without performing any prior bacterial culturing and fluorescence labeling. First, we evaluated the applicability of Glycan-seq for bacterial profiling using the cultured representatives of gram-positive and gram-negative bacteria. We then used Glycan-seq to analyze the glycome alteration on the gut microbiota of young and old mice. Further, 16S rRNA gene sequencing was performed to analyze the differences in the bacterial composition of the gut microbiotas in young and old mice.

## Methods

### Microbial culture

*Escherichia coli* (Migula) (ATCC 700926) was cultured overnight at 37°C in M9 culture medium, whereas *Deinococcus radiodurans* (ATCC BAA-816) was also cultured overnight at 30°C in TGY medium. The abundance and size of cells were analyzed using a particle counter (CDA 1000; Sysmex Corporation, Hyogo, Japan).

### Mice

Young (14–20 days old) and old (12 months old) C57BL/6J mice were used in this study. The mice were derived or purchased from Charles River Laboratories and Japan SLC (Shizuoka, Japan). Male mice were used for all the experiments. The mice were housed under specific pathogen-free conditions in the Laboratory Animal Resource Center at the University of Tsukuba, Japan.

### Fecal sample collection and microbiota isolation

Mice were placed inside an autoclaved cage for 30–60 min, and the excreted feces were collected using sterilized forceps. The collected feces were frozen at –20°C until use. The mouse fecal microbiota was isolated using the density gradient method [16]. Briefly, approximately 20 mg of feces was homogenized in 0.5 ml phosphate-buffered saline (PBS) at 4°C by shaking at 750 rpm overnight. After homogenization, the supernatant was collected and transferred to the top of a Nycodenz solution (80% w/v in water; Cosmo Bio Co., Ltd., Tokyo, Japan). The solution was then centrifuged at 10,000 × *g* for 40 min at 4°C. The middle layer containing the microbiota was collected and further washed with PBS. The numbers and sizes of bacterial cells from each sample were quantified using a particle counter (CDA-1000; Sysmex Corporation).

### Preparation of DNA-barcoded lectins

Lectins were conjugated to the DNA oligonucleotide as previously described [15]. Briefly, 100 μg of each lectin was dissolved in 100 μl of PBS mixed with dibenzocyclooctyne-N-hydroxysuccinimidyl ester (DBCO-NHS) (Funakoshi Co., Ltd., Tokyo, Japan) at 10 times the molar amount and then incubated in the dark for 1 hour at 20°C. DBCO-NHS was inactivated by adding 10 μl of 1 M Tris and incubating the mixture in the dark for 15 min at 20°C. The excess DBCO-NHS was removed using Sephadex G-25 desalting columns (GE Healthcare Japan Co., Tokyo, Japan). The DBCO-labeled lectin product (100 μg/mL) was mixed with 5’-azide-modified DNA oligonucleotides (Integrated DNA Technologies, KK, Tokyo, Japan) at 10 times the molar amount. The conjugated lectin-DNA oligonucleotide was purified by removing unbound nucleotides and selecting only the lectins with the glycan-binding affinity, which was achieved by affinity chromatography using the appropriate sugar-immobilized Sepharose 4B-CL (GE) based on the glycan-binding specificity of each lectin.

### Glycan-seq

Bacterial cells (1 × 10^7^) were suspended in PBS containing 1% bovine serum albumin (BSA) and incubated with 39 DNA-barcoded lectins at a final concentration of 0.5 μgml^-1^ at 4°C for 1 h. The cells were washed three times with 1 ml of PBS/BSA to liberate oligonucleotides after which it was diluted ten times (1 × 10^6^) and then were UV-irradiated at 365 nm, 15 W, for 15 min using a UVP Blak-Ray XX-15L UV Bench Lamp (Analytik Jena, Kanagawa, Japan). The liberated oligonucleotides were then amplified using NEBNext Ultra II Q5 (New England BioLabs Japan Inc., Tokyo, Japan), i5-index, and i7-index primers containing cell oligonucleotide sequences. PCR reactions were performed as follows: 1 cycle of denaturation for 45 sec at 98°C; 20 cycles of denaturation for 10 sec at 98°C, followed by 50 sec at 65°C; and 1 cycle of extension for 5 min at 65°C. The PCR products were then purified using the Agencourt AMPure XP Kit (Beckman Coulter, Inc., Tokyo, Japan) following the manufacturer’s protocol. The size and quantity of the PCR products were analyzed using MultiNA (Shimadzu Co., Kyoto, Japan). The PCR products (4 nM from every sample) were treated with the MiSeq Reagent Kit v2 (50 cycle format; Illumina KK, Tokyo, Japan) and sequenced using the MiSeq Sequencer (26 bp, paired-end) (Illumina KK, Tokyo, Japan).

### Analysis of Glycan-seq data

The DNA barcodes derived from lectins were directly extracted from the reads in FASTQ format. The number of DNA barcodes bound to each cell was counted using a barcode DNA counting system (Mizuho Information & Research Institute, Inc., Tokyo, Japan) [15]. The first three bases in each read were removed to better match the DNA barcode sequence. In cases of mismatch, we allowed a maximum of two mismatches in the flanking region and one mismatch in the middle region. The total number of each of the DNA barcodes was divided by the total number of lectin barcodes and expressed as a percentage (%) for each lectin. Statistically significant levels of lectins in the Glycan-seq were evaluated by *t*-tests, setting the levels of significance at *P* < 0.01 for cultured bacteria and *P* < 0.05 for the young and old gut microbiota.

### Flow cytometry analysis

Approximately 1 × 10^7^ cells of *E. coli* and *D. radiodurans* were incubated with 10 μg of R-phycoerythrin-conjugated lectins for 1 hour on ice. BSA-conjugated lectin was used as a negative control. Flow cytometry data were acquired on a CytoFLEX System (Beckman Coulter, Inc., Brea, CA) and analyzed using the FlowJo software v10.6 (BD, Franklin Lakes, NJ).

### Microbial DNA extraction from mouse feces

Genomic DNA was isolated from the microbial fraction collected from mouse feces (as described above) by a bead-beating method implemented using the ISOSPIN Fecal DNA Kit (Nippon Gene Co., Ltd, Japan). The isolated DNA was eluted in 50 μl TE buffer (pH 8.0) provided in the kit.

### 16S rRNA gene sequencing

Sequencing libraries were prepared from the V3–V4 hypervariable region of 16S rRNA gene, following the protocol entitled “16S Metagenomic Sequencing Library Preparation” from Illumina [17]. The V3–V4 hypervariable region of 16S rRNA gene was amplified using the following primers: forward: 5’-TCGTCGGCAGCGTCAGATGTGTATAAGAGACAGCCTACGGGNGGCWGCAG-3’; reverse: 5’-GTCTCGTGGGCTCGGAGATGTGTATAAGAGACAGGACTACHVGGGTATCTAATCC-3’). The 25 μl PCR reaction was performed using a KAPA HiFi HotStart ReadyMix (Roche) and contained 1 μl of extracted fecal microbial DNA and 1 μM of each primer. The reaction cycles consisted of initial denaturation at 98°C for 2 min; followed by 25 cycles of denaturation at 98°C for 15 sec, annealing at 56°C for 30 sec, and elongation at 72°C for 30 sec; and a final elongation at 72°C for 5 min.

Next, a second PCR was performed using Illumina index primers and the following reaction cycle: initial denaturation at 95°C for 3 min; followed by 8 cycles of denaturation at 95 °C for 30 sec, annealing at 55°C for 30 sec and elongation at 72°C for 30 sec; and a final elongation at 72°C for 5 min. The amplicons were quantified using MultiNA (Shimadzu, Japan), a microchip electrophoresis system for DNA/RNA analysis. The amplicons were sequenced using the Illumina MiSeq 2 × 250 bp platform with a MiSeq Reagent Nano Kit V2 (Illumina).

### 16S rRNA gene sequence analysis

The raw sequence reads were analyzed using QIIME2 (2020.8) [18]. The reads were first demultiplexed; then, the DADA2 [19] plugin was used for quality control, read trimming, and assembly. Trimming took into consideration the information needed to merge the paired reads. Amplicon sequence variants (ASVs) were generated by DADA2 analysis, which were then classified to family and genus levels using the q2-feature-classifier [20], a Naïve Bayes machine learning classifier plugin in the QIIME2. Operational taxonomic units (OTUs) were generated by the RESCRIPt QIIME2 plugin running a feature classifier trained on the V3–V4 region of the 16S rRNA gene using a preformatted SILVA 138 reference database [21, 22]. An equal sampling depth of 10,000 was selected for every sample for assessing the diversities. α-diversity was measured by Faith’s phylogenetic diversity (PD) metrics, and significance (*p* < 0.05) was statistically calculated using Kruskal-Wallis (pairwise) analysis. Using principal coordinate analysis (PCoA) from the UniFrac metrics analysis, β-diversity was calculated [23].

### Microbe isolation by lectin

SSA and TJAI lectins were labeled with biotin and used at a concentration of 1 μg/ul. The labeled lectins were incubated with streptavidin-conjugated Dynabeads (Thermo Fisher, Waltham, Massachusetts, USA) in a shaker set at 1,400 rpm at 4°C for 1 hour. The conjugated beads were washed; then, 2 × 10^7^ microbial cells from young and old mice samples were incubated with the beads in a shaker at 700 rpm at 4°C overnight. The bound microbes were isolated using a magnetic stand and eluted with 2M lactose.

### Correlation analysis

The x of lectins and the y of microbial communities were plotted using the corrplot package in R. The analysis yielded Spearman correlation coefficients evaluated at *p* < 0.05.

## Results

### Glycomic profiling of the gut microbiota by Glycan-seq

We aimed to develop a strategy to profile the glycome of the intact gut microbiota without prior fluorescence labeling using Glycan-seq [15]. Cultured bacterial cells were incubated with DNA-barcoded lectins, which, upon binding, released their DNA barcodes after UV exposure, because lectins were conjugated with DNA barcodes via a photocleavable linker (Fig. 1). The lectins used in this study cover a wide range of glycan structures, including sialylated, galactosylated, GlcNAcylated, mannosylated, and fucosylated glycans (Table S1). The released DNA barcodes were recovered, amplified, and analyzed by next-generation sequencing. The number of each DNA barcode was divided by the total number of lectin barcodes and expressed as percentage (%) values for each lectin. The microbiotas obtained from the young and old mice were also analyzed by 16S rRNA gene sequencing to identify the populations of bacteria.

**Fig. 1.**
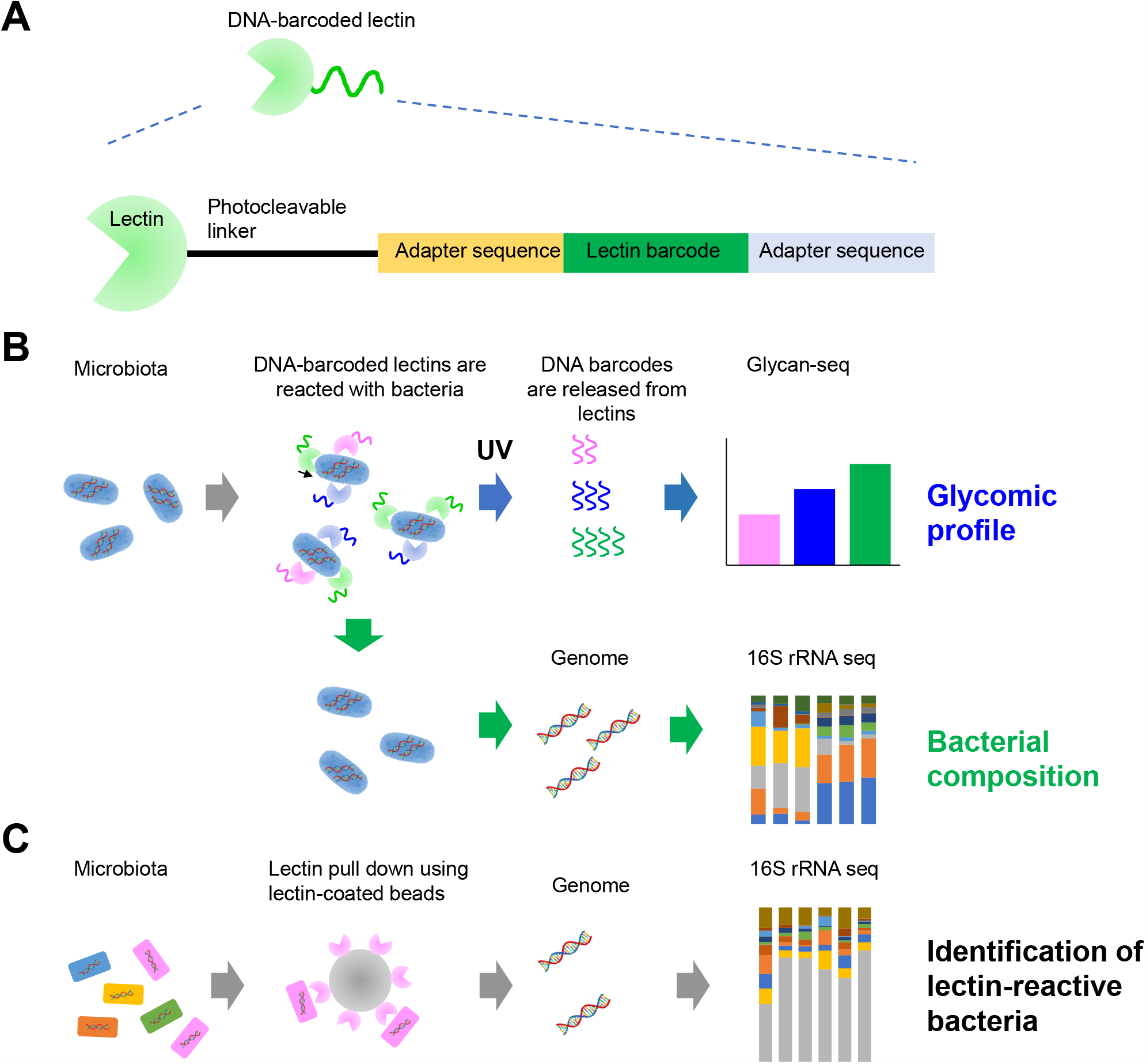
Glycomic profiling of microbiota by Glycan-seq. (A) Schematic representation of a DNA-barcoded lectin. (B) Experimental workflow of the Glycan-seq and 16S rRNA sequencing of microbiota. (C) Schematic representation of lectin pull-down followed by 16S rRNA sequencing to identify lectin-reactive bacteria.

### Glycan-seq of the cultured bacteria

We first evaluated whether Glycan-seq can be used to profile the glycans of cultured gram-positive *D. radiodurans* and gram-negative *E. coli* (Fig. 2, Table S2). Bacterial cells (1 × 10^7^) were incubated with DNA-barcoded lectins, and the DNA barcodes that were released from 1 × 10^6^ bacterial cells by UV irradiation were counted by sequencing. The resulting lectin binding signals were first analyzed by the hierarchical cluster analysis (Fig. 2A). The two types of bacteria were clearly separated into two different clusters, where the *x*-axis shows the lectins used and the *y*-axis shows the bacterial species. Several lectins differentially bound to *D. radiodurans* and *E. coli*. Specifically, mannose-binders (rGRFT and rBanana) reacted at significantly higher levels to *D. radiodurans* (*p* < 0.01), whereas GalNAc (HPA)-, Galβ1-3GlcNAc/GlcNAc (rABA, rSRL)-, GlcNAc (rPVL)-, and rhamnose (CSA)-binders exclusively reacted with *D. radiodurans* (*p* < 0.01, Fig. 2B).

**Fig. 2.**
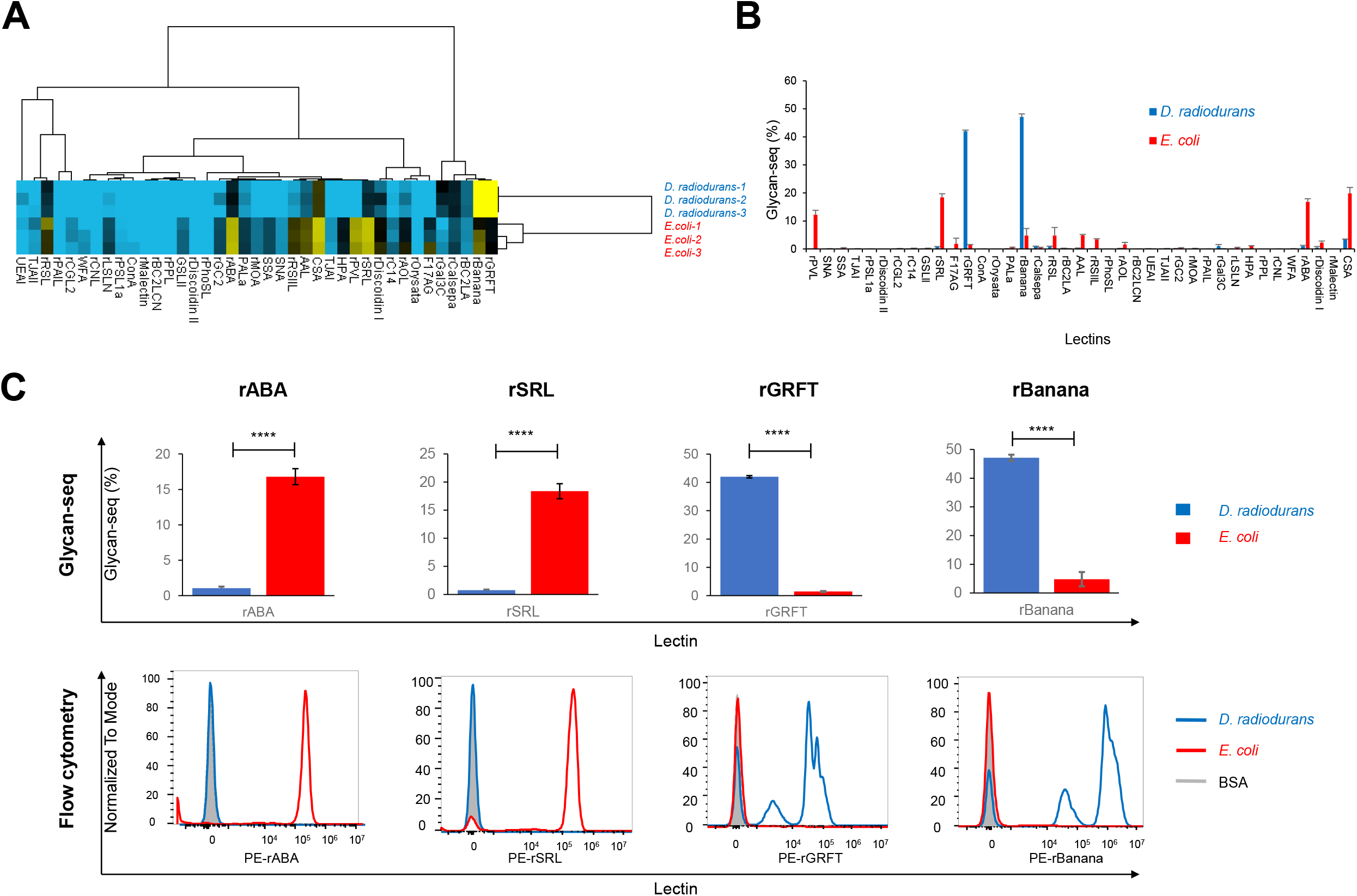
Glycomic profiling of the cultured bacteria by Glycan-seq. (A) Hierarchical cluster analysis of *D. radiodurans* and *E. coli* using Glycan-seq data. (B) Graphical representation of Glycan-seq intensity data of *D. radiodurans* and *E. coli*. (C) Comparison between Glycan-seq (top panel) and flow cytometry data (bottom panel). Blue: *D. radiodurans*; red: *E. coli*.

We validated the results of lectin binding to *D. radiodurans* and *E. coli* obtained by Glycan-seq using flow cytometry analysis, which is considered the gold standard. Lectins that specifically bound to the two different types of bacteria were identified by *t*-test analysis (Table 1). Sixteen lectins specifically bound at significant levels (*p* < 0.01, Fig 2B). Based on the signal intensity (average intensity for positive cells, >0.5) obtained from Glycan-seq and the *t*-ratio (>20) from statistical analysis, we selected the following four lectins for flow cytometry analysis (Fig. 2B and Table 1): mannose-binders (rGRFT, rBanana) that generated higher signals in *D. radiodurans* and Galβ1-3GalNAc/GlcNAc-binders (rABA, rSRL) that generated higher signals in *E. coli*.

**Table 1.** Lectins with significantly different signals between Gram-positive and -negative bacteria selected by Glycan-seq

| Lectins | Rough specificity | *t*-ratio | *P*-value | Average intensity of *D. radiodurans* | Average intensity of *E. coli* |
| --- | --- | --- | --- | --- | --- |
| **rGRFT<sup>1</sup>** | **Man** | **163.19** | **0.000000338** | **41.99** | **1.44** |
| SSA | α2-6Sia | 124.51 | 0.000000973 | 0.06 | 0.47 |
| rMOA | αGal (B) | 88.106 | 0.00000378 | 0.03 | 0.23 |
| SNA | α2-6Sia | 42.021 | 0.0000709 | 0.02 | 0.13 |
| **rBanana** | **Manα1-2Manα1-3(6)Man** | **27.118** | **0.0003958** | **47.13** | **4.81** |
| AAL | α1-2Fuc (H), α1-3Fuc (Lex),<br>α1-4Fuc (Lea) | 26.435 | 0.0004259 | 0.31 | 4.89 |
| **rABA** | **Galβ1-3GalNAc (T),**<br>**GlcNAc** | **23.592** | **0.0006505** | **1.05** | **16.79** |
| **rSRL** | **Galβ1-3GalNAc (T),**<br>**GlcNAc** | **22.718** | **0.0007336** | **0.75** | **18.38** |
| rRSIIL | α1-2Fuc (H), α1-3Fuc (Lex),<br>α1-3Fuc (Lea) | 21.691 | 0.0008548 | 0.11 | 3.42 |
| HPA | αGalNAc (A, Tn) | 18.832 | 0.0014504 | 0.03 | 1.09 |
| rPALa | Man5, biantenna | 17.279 | 0.001973 | 0.13 | 0.62 |
| rPPL | $\alpha$,βGalNAc (A, Tn,<br>LacDiNAc) | 15.275 | 0.0031018 | 0.00 | 0.01 |
| rPVL | Sia, GlcNAc | 13.279 | 0.0051914 | 0.04 | 12.23 |
| CSA | Rhamnose, Gala1-4Gal | 13.216 | 0.0051914 | 3.52 | 19.80 |
| TJAI | α2-6Sia | 12.161 | 0.0067989 | 0.01 | 0.12 |
| rBC2LA | αMan, High-man | 11.036 | 0.0095384 | 0.27 | 0.15 |

<sup>1</sup> Lectins in bold characters were selected for further flow cytometry analysis

Flow cytometry using fluorescently labeled mannose-binders (rGRFT, rBanana) generated a higher peak signal in *D. radiodurans*, whereas similar analysis using Galβ1-3GalNAc/GlcNAc-binders (rABA, rSRL) generated a higher peak signal in *E. coli* (Fig. 2C). Thus, the results of flow cytometry agreed with those obtained by Glycan-seq (Fig. 2B). Taken together, these results indicate that bacterial glycan profiles generated by Glycan-seq can be recapitulated by flow cytometry analysis.

### Glycan-seq of the gut microbiota from young and old mice

As Glycan-seq was applicable for both gram-positive and gram-negative bacteria, we used this approach to profile the gut microbiota. Bacterial cells were from the fecal microbiota of young, preweaned (14–20 days) and old (12 months) mice (*n* = 3), and the numbers and sizes of cells are shown in Table 2. On average, the bacterial cells from the feces of young mice numbered 1.6 × 10^10^ cells/g with an average diameter of 1.2 μm, whereas those from old mice numbered 1.8 × 10^10^ cells/g with an average diameter of 0.97 μm. The fecal microbiotas (1 × 10^7^ cells) of young and old mice were then subjected to Glycan-seq, followed by hierarchical cluster analysis. Figure 3A shows that the gut microbiotas of young and old mice are separated into two clusters based on Glycan-seq (Table S3), demonstrating that the gut microbiotas of young and old mice have distinct glycan profiles. Statistical analysis on the lectin intensity data obtained from Glycan-seq reveals four lectins that significantly differentiated the young and old microbiotas (Table 3). These lectins are α2-6Sia-binders (SSA, TJAI) and Galβ1-3GalNAc/GlcNAc-binders (rSRL, rABA) (Fig. 3B), all of which were detected at higher levels in the young microbiota. Previous studies have shown that the composition and diversity of the gut microbiota change with age [24]. Notably, the levels of some lectins, including rhamnose (CSA) and fucose-binders (rPhoSL, rBC2LCN) (Fig. 3B), tend to be higher in old microbiota. However, the difference is not statistically significant (Fig. 3B), which may be due to the increases in microbial diversity and mouse-to-mouse variation in the composition of microbiota in old mice. Nevertheless, our data suggest that our newly developed Glycan-seq technology successfully profiled the glycome of the gut microbiota.

**Table 2.**
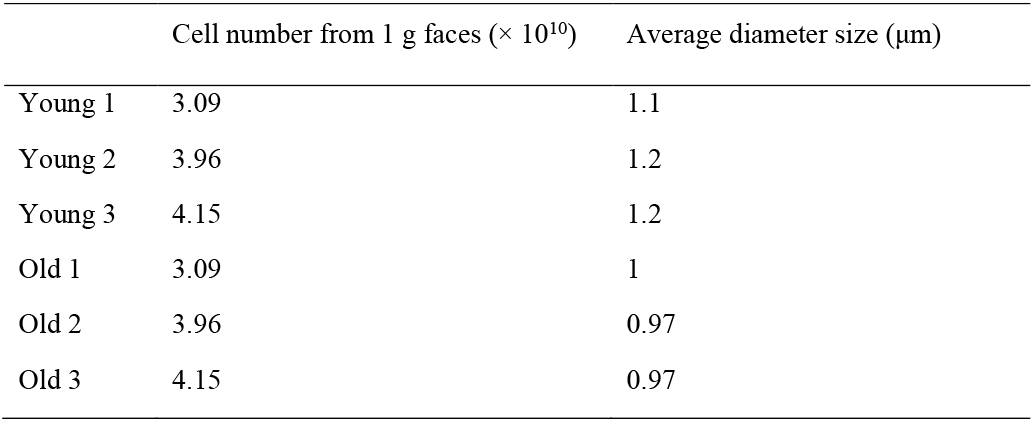
Total number of bacterial cells and the average size prepared from the microbiotas of young and old mice

**Fig. 3.**
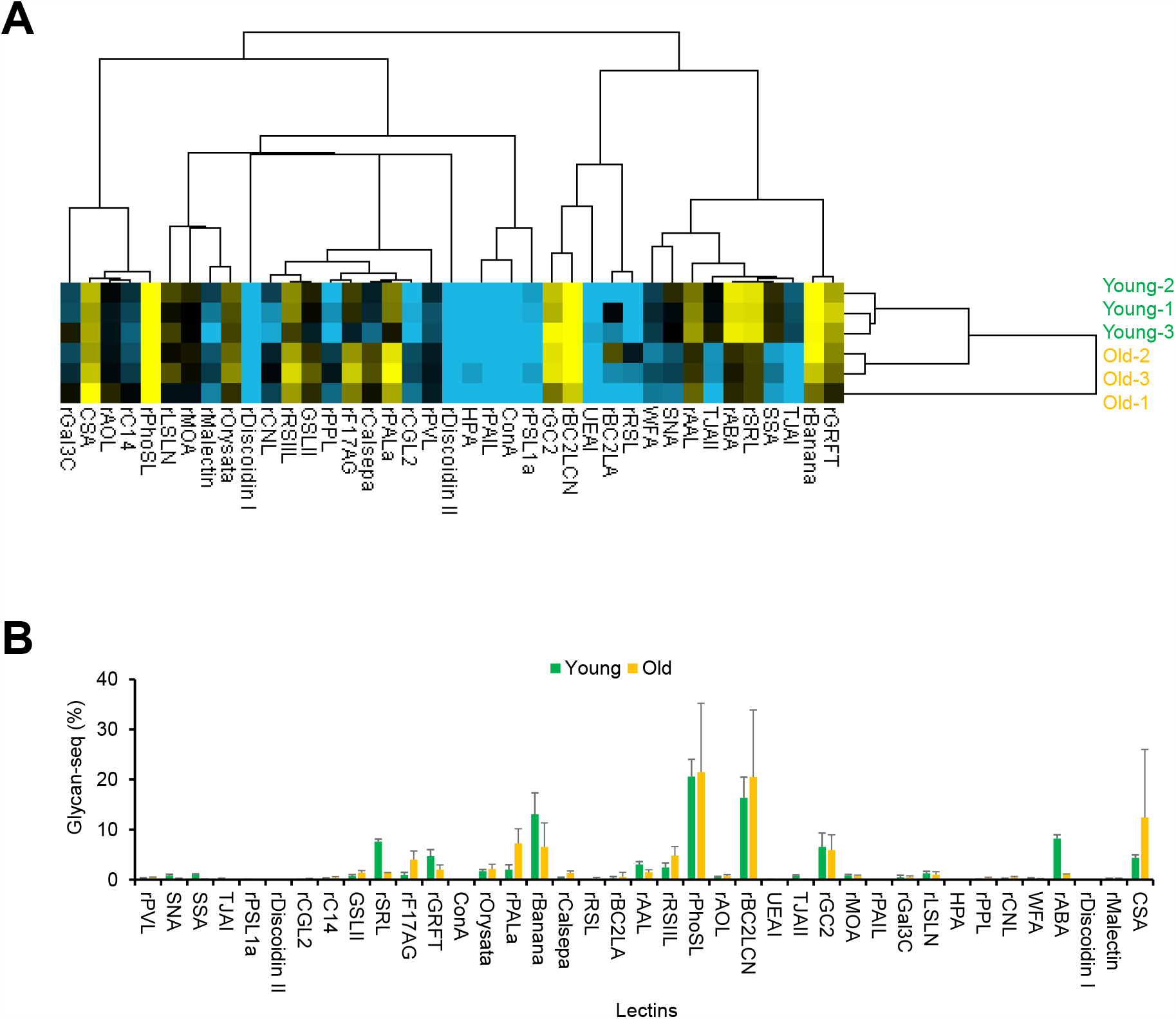
Glycomic profiling of the gut microbiota of young and old mice. (A) Hierarchical clustering analysis of the gut microbiota of young and old mice using Glycan-seq data. (B) Graphical representation of the Glycan-seq data of the gut microbiota of young and old mice.

**Table 3.**
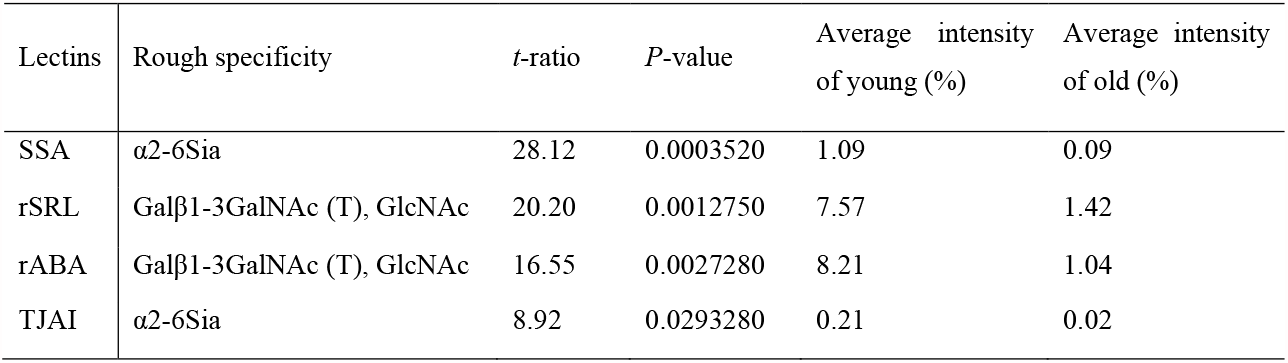
Lectins with significantly different signals between young and old microbiota selected by Glycan-seq

### Differences in the composition of the gut microbiota of young and old mice

Based on 16S rRNA sequencing, the α- and β-diversity of young and old gut microbiota differed (Figs. 4A and 4B). This result is similar to that of a previous study [25] that reported differing compositions of the gut microbiota of young and old mice. The analysis of the ASVs of the metagenomic data of 16S rRNA gene sequences using QIIME2 [18] shows family-level differences between young and old microbiotas. Specifically, the relative abundance levels of *Lactobacillaceae, Enterobacteriaceae, Pseudomonadaceae*, and *Gemellaceae* were higher in young microbiota. In contrast, *Rikenellaceae, Erysipelotrichaceae, Muribaculaceae, Bifidobacteriaceae*, and *Lachnospiraceae* were higher in old microbiota (Fig. 4C). Therefore, these differences in gut microbiota diversity may explain the different glycan profiles of young and old microbiotas, as revealed by Glycan-seq analysis.

**Fig. 4.**
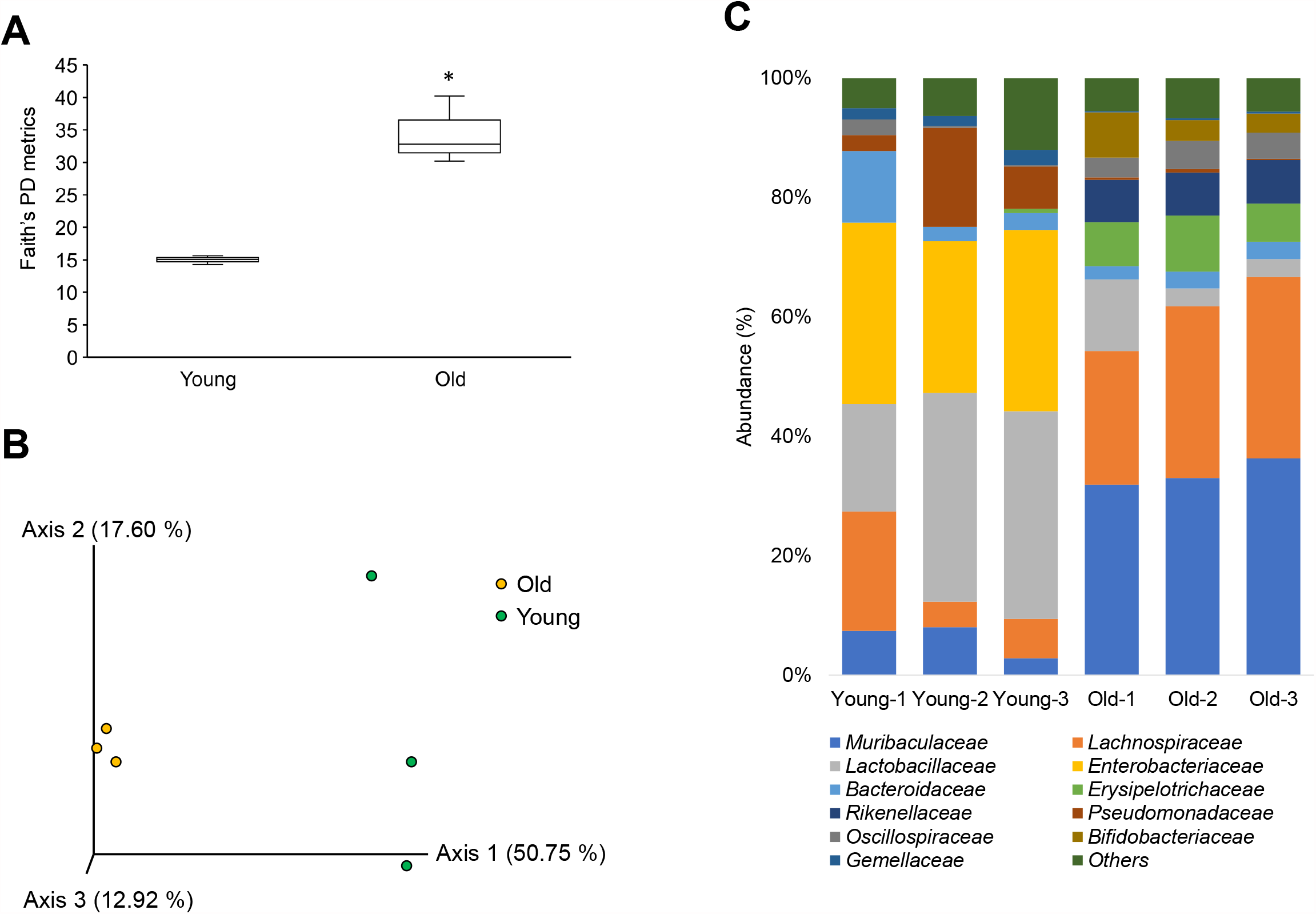
Comparison of the bacterial abundance between the young and old mice microbiota. (A) Boxplot of Faith’s phylogenetic diversity (PD) metrics analysis for the α-diversity. Statistical significance (*p* <0.05) is denoted with an asterisk (*) calculated by Kruskal-Wallis pairwise analysis. (B) β-diversity analysis by principal coordinates analysis (PCoA) of unweighted UniFrac distance. (C) Stacked bar plot showing the taxonomy of the differential bacterial abundance in the young and old mice microbiota obtained from each mouse by 16S rRNA sequencing (*n* = 3 for each age group). Each colored bar plot indicates the family of bacteria identified, and for clarity, only the most abundant 11 families are shown, and the remaining are shown as others

### Changes in gut microbiota diversity are associated with the different glycans detected in the gut microbiota

We investigated whether the differences in the relative abundance levels of bacterial families are correlated with the lectins identified by Glycan-seq analysis through Spearman’s correlation analysis. The analysis revealed the correlation between the relative abundance values of the bacteria and the lectin signal intensities of the young and old microbiotas. Lectins with variable correlations to the bacterial abundance were also identified (Fig. 5). α2-6Sia-binders (SSA, TJAI) and Galβ1-3GalNAc/GlcNAc-binders (rSRL, rABA) had significantly higher intensities in the young microbiota (Fig. 3). Moreover, the signal intensities of α2-6Sia-binders (SSA, TJAI) were positively correlated with those of Galβ1-3GalNAc/GlcNAc-binders (rSRL, rABA)(Fig. 5). These lectins were positively correlated with levels of *Lactobacillaceae, Enterobacteriaceae*, and *Gemilaceae* but were negatively correlated with the *Muribaculaceae, Lachnospiraceae, Erysipelotrichaceae*, and *Rikenellaceae*.

**Fig. 5.**
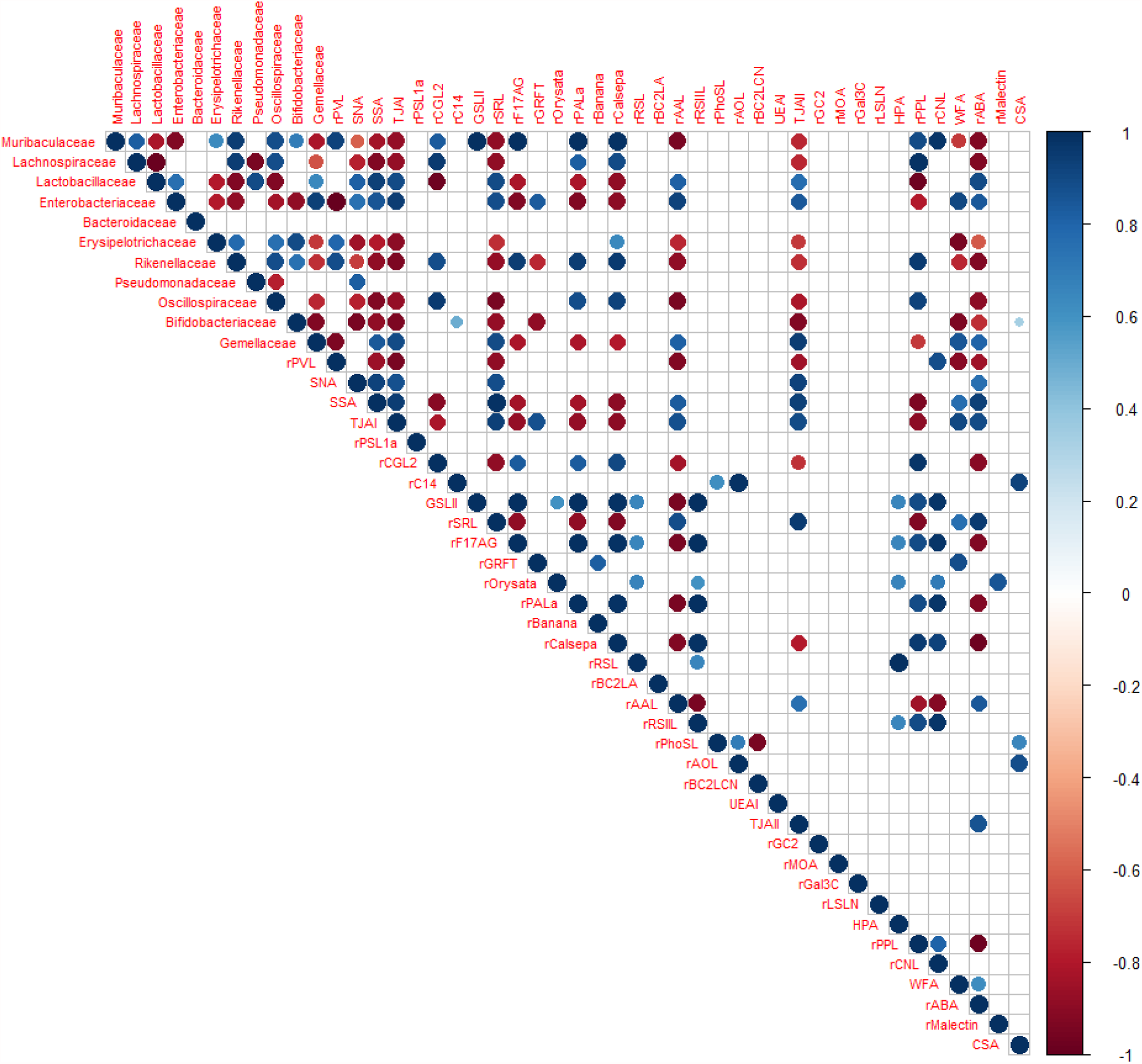
Correlation between bacterial family and lectin. Spearman correlations between the abundant bacterial family from 16S rRNA sequencing and the lectin intensity from the Glycan-seq. The correlations represented were statistically significant (p < 0.05), and the circle’s size represents the strength of correlations. Red: negative; blue: positive correlations.

### Identification of sialylated bacteria in the gut microbiota of young mice

The signal intensities of α2-6Sia-binders (SSA, TJAI) were significantly higher in the microbiota of young mice. We were thus interested in investigating which bacteria are covered with Sia, a monosaccharide that is primarily found at the nonreducing end of glycoconjugates in eukaryotes and is involved in a variety of cell–cell interactions and cell–molecule recognition [26]. Several species of pathogenic bacteria display Sia on their outer surfaces, which masks them from the host immune system [27]. We used a lectin pull-down assay to determine which bacteria in the microbiota of young mice react with α2-6Sia-binders. The assay involved the incubation of bacterial cells (isolated from young and old mice) together with magnetic beads coated with α2-6Sia-binders (SSA, TJAI). The average number of cells recovered from the young microbiota by the assay was 2.5 × 10^6^ cells/g, with an average diameter of 1.1 μm, whereas only approximately one-hundredth of bacterial cells (an average of 1.9 × 10^4^ cells/g with an average diameter of 0.6 μm) were obtained in the old mice (Table 4). The incubation of both types of microbiotas with lectin-coated beads resulted in the recovery of more cells from the young microbiota, indicating that more bacterial cells with glycans that react with α2-6Sia-binders are present in young mice (Fig. 6).

**Table 4.** Cell number and average size of bacteria pull down by SSA- and TJAI-coated beads

|  | SSA pull down |  | TJAI pull down |  |
| --- | --- | --- | --- | --- |
| | Cell number<br>($\times 10^4$) from 1 g<br>feces | Average diameter<br>size ($\mu\text{m}$) | Cell number<br>($\times 10^4$) from 1 g<br>feces | Average diameter<br>size ($\mu\text{m}$) |
| Young 1 | 263 | 1.2 | 217 | 1.1 |
| Young 2 | 186 | 1.1 | 285 | 1.09 |
| Young 3 | 222 | 1.09 | 286 | 1.1 |
| Old 1 | 2.63 | 0.61 | 2.31 | 0.65 |
| Old 2 | 1.96 | 0.58 | 2.9 | 0.6 |
| Old 3 | 0.82 | 0.62 | 1.2 | 0.61 |

**Fig. 6.**
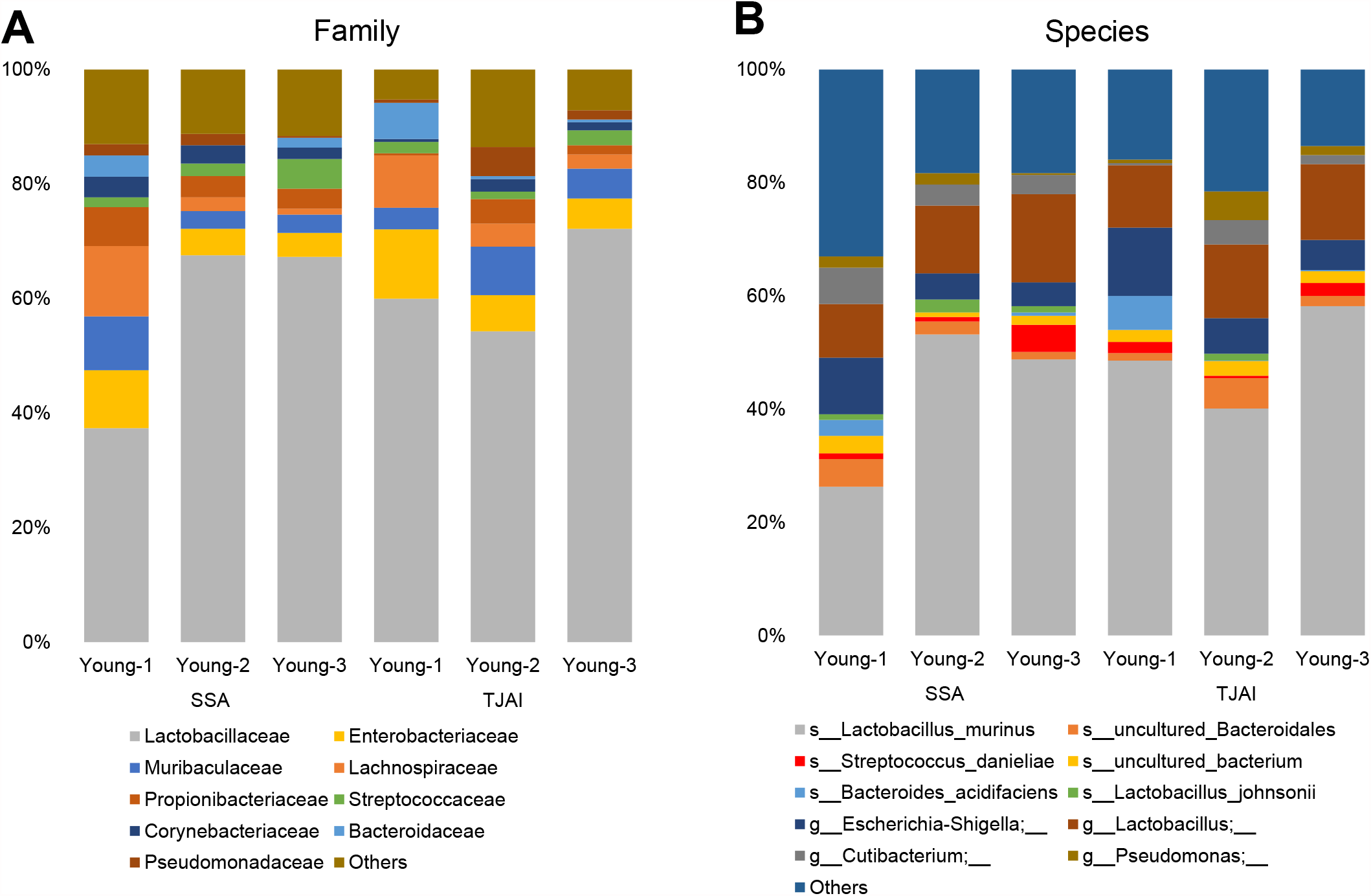
Family of bacteria reactive to α2-6Sia identified by lectin pull-down and 16S rRNA sequencing from the young microbiota. Stacked taxa bar plot represents the (A) family (B) species of bacteria pull-downs by the SSA and TJAI, α2-6Sia binders. Each colored bar plot indicates the family or genus of bacteria identified, and for clarity, only the most abundant families or species are shown, and the remaining are shown as others.

The identification of the taxonomic groups of bacteria pulled down by the α2-6Sia-binders is based on their 16S rRNA gene sequences. The ASV analysis (using QIIME2) of the 16S rRNA gene sequences from the metagenomic data shows the phylogenetic distribution and relative abundance of bacteria pulled down by SSA and TJAI. The results identify the families of bacteria present at higher levels in the young microbiota (Fig. 6). Both lectins detected mainly *Lactobacillaceae, Lachnospiraceae, Enterobacteriaceae*, and *Muribaculaceae* (Fig. 6). At the genus level, these bacteria consisted mainly of *Lactobacillus, Pseudomonas, Escherichia-Shigella*, and *Streptococcus*. Therefore, these bacterial families and genera are likely modified by sialylated glycans. These results show that the bacteria recovered from the gut microbiota using lectins are those covered by Sia.

## Discussion

The genome and metabolome are the two major omics data acquired for the microbiota. In this sense, the information of the intact bacterial cell surface molecules without prior *in vitro* culturing, which directly mediate microbe–host interactions, coundn’t be acquired. Actually, bacterial cell surfaces are coated with a diverse array of glycans that play pivotal roles in various biological processes. In particular, they mediate microbe–host interactions during the onset and development of infectious diseases and symbiotic interactions. However, our understanding of the glycome of the gut microbiota remains limited because of the lack of appropriate methods of analysis. In this study, we have developed a new sequencing-based technology that can analyze the glycome of bacteria in an intact form. The Glycan-seq technology offers several advantages: (1) Live bacterial cells can be analyzed without the need for prior culturing. (2) Fluorescence labeling of bacteria is not required. (3) A relatively low number of bacterial cells (∼10^6^ cells) are required for the analysis. (4) Glycomic profiles can be acquired using a conventional next-generation sequencer, the same instrument used for 16S rRNA gene sequencing.

The diversity of the gut microbiota of old mice differed from that of young mice. The relative abundance levels of *Lactobacillaceae, Enterobacteriaceae, Pseudomonadaceae*, and *Gemellaceae* were higher in young than in old microbiota, whereas those of *Rikenellaceae, Erysipelotrichaceae, Muribaculaceae, Bifidobacteriaceae*, and *Lachnospiraceae* were higher in old than in young microbiota. These differences in the gut microbiota diversity may be due to host differences in feeding habits; young mice are nursed on mother’s milk, whereas old mice are fed with normal chow diets. In humans, breastfeeding babies have more *Lactobacilli* in their gut microbiota than those of milk formula-fed babies [28]. These findings and the results of this study (Fig. 4) suggest the common presence of *Lactobacilli* in the gut of breastfed animals. A study on calorie-restricted mice found reduced levels of *Lactobacillus*, which was negatively correlated with mice lifespan [29]. Furthermore, the abundance of *Lactobacillus murinus* is higher in calorie-restricted mice, and this species promoted anti-inflammation, which may play an important role during aging [30].

Several studies have sought to understand how the gut microbiota changes during aging and the biological significance of such changes [31]. The results from longitudinal studies on fecal samples from various individuals of different ages show age-related changes in the diversity and composition of the human gut microbiota [32, 33]. The composition of the gut microbiota of older adults is unique, and the α-diversity of this microbiota increases with age, suggesting a correlation between the composition of the gut microbiota and physiological aging [33].

In this study, we report for the first time that the glycome of the gut microbiota changes during aging. Glycan-seq technology was able to profile the microbial glycomes of young and old mice, and interestingly, α2-6Sia-binders reacted at significantly higher levels with the young microbiota, suggesting that the levels of sialylated bacteria decrease during aging. The bacterial families that reacted most with α2-6Sia-binders are *Lactobacillaceae, Lachnospiraceae, Enterobacteriaceae*, and *Muribaculaceae*. Previous findings that some *Lactobacillus* species express genes involved in the catabolism of Sia [27] are consistent with our results. The presence of Sia in *Lactobacillus* species has also been previously reported [34]. Several pathogenic bacteria such as enterohemorrhagic *Escherichia coli, Haemophilus influenzae, H. ducreyi, Pasteurella multocida, Neisseria gonorrhoeae, N. meningitidis, Campylobacter jejuni*, and *Streptococcus agalactiae* are well known to display Sia on their outer surfaces, which masks them from the host immune system. These bacteria have developed different mechanisms for obtaining Sia, including the *de novo* biosynthesis of Sia (*E. coli, N. meningitidis*), Sia scavenging (*N. gonorrhoeae*), and precursor scavenging (*H. influenzae*) [27]. One of the functions of Sia is the regulation of innate immunity by providing a mechanism of identifying self from nonself. However, various microbes have evolved a counter-mechanism that works by decorating the bacterial cell surface with similar Sia modifications [35]. Sia that decorates the bacterial surface regulates the host immune system by interacting with sialic acid-binding immunoglobulin-type lectins (Siglecs) [36]. The presence of the Sia on the surface of bacteria also protects them against invading bacteriophage by blocking the relevant underlying receptors [37]. Therefore, the presence of Sia on the surface of bacteria in the gut microbiota of young mice suggests that these microbes are protected from the host’s innate immune surveillance system and from bacteriophage, and their establishment proceeds from an initial colonization by microbes in the gut of young mice [35].

Most bacteria obtain Sia by scavenging it from the surrounding environment [27]. Therefore, the glycans of bacteria cultured *in vitro* most probably differs from that of the same bacteria growing in the gut. Given this situation, Glycan-seq is useful because it can capture the glycomic information of gut bacteria *in situ*, because it does not require prior culturing *in vitro*.

The following are the limitations of the current method: (1) the glycomic profile of single bacterial cells cannot be obtained; and (2) the method is unable to determine the detailed structure of glycans. We aim to solve the first limitation by developing a method of simultaneously analyzing the glycome and genome of single bacterial cells.

## Conclusions

Our results suggest that the Glycan-seq method is an excellent choice for profiling the glycome of the gut microbiota. Our data provides important (and previously unknown) details about the changes in the glycome of the gut microbiota during aging. Glycan-seq analysis, in parallel with 16S rRNA gene sequencing, can identify the bacteria modified with Sia. It will be interesting to apply the Glycan-seq technology in future studies, seeking to understand how the glycome of the gut microbiota changes in response to dietary changes and disease development. Moreover, application of the Glycan-seq method to profile the glycome of a single bacterial cell, along with the bacterial identification, will help researchers understand the glycome architecture of the gut microbiota and its interaction with the host. In addition, our technology can also be applied to profile the glycomes of other bacterial communities, such as those in the soil, deep ocean, and volcanic deposits.

## Acknowledgments

The authors would like to thank Professor Hiromi Yanagisawa and the Animal Resource Center (ARC) staff of the University of Tsukuba for assisting in collecting the mouse feces; Drs. Jonguk Park and Jun Kunisawa at National Institutes of Biomedical Innovation, Health and Nutrition, and Drs. Hiroyuki Kusada and Hideyuki Tamaki at the National Institutes of Advanced Industrial and Technology for technical advice on 16S rRNA sequencing; Ms. Sunanda Keisham for her helpful discussion during the experiment.

## Authors’ contributions

LO performed experiments and data analysis and wrote the paper. FM performed experiments and data analysis. HT designed this study, led this study, and wrote the paper. All authors provided feedback and contributed to the research and final manuscript.

## Availability of data and materials

All the data generated and analyzed have been included in the article or as supplementary tables and files. The raw 16S rRNA amplicon sequencing data are deposited and publicly available from European Nucleotide Archive (ENA) at EMBL-EBI under accession number PRJEB45936 (https://www.ebi.ac.uk/ena/browser/view/PRJEB45936).

## Declarations

### Ethics approval and consent to participate

Not applicable

### Consent for publication

Not applicable

## References

1. Costello EK, Lauber CL, Hamady M, Fierer N, Gordon JI, Knight R. Bacterial Community Variation in human body habitats across space and time. Science (80-). 2009;326:1694LP–1697. doi:10.1126/science.1177486.

2. Qin J, Li R, Raes J, Arumugam M, Burgdorf KS, Manichanh C, et al. A human gut microbial gene catalogue established by metagenomic sequencing. Nature. 2010;464:59–65. doi:10.1038/nature08821.

3. Eckburg PB, Bik EM, Bernstein CN, Purdom E, Dethlefsen L, Sargent M, et al. Diversity of the human intestinal microbial flora. Science (80-). 2005;308:1635LP–1638. doi:10.1126/science.1110591.

4. Valdes AM, Walter J, Segal E, Spector TD. Role of the gut microbiota in nutrition and health. BMJ. 2018;361:k2179. doi:10.1136/bmj.k2179.

5. Kayama H, Okumura R, Takeda K. Interaction between the microbiota, epithelia, and immune cells in the intestine. Annu Rev Immunol. 2020;38:23–48. doi:10.1146/annurev-immunol-070119-115104.

6. Carabotti M, Scirocco A, Maselli MA, Severi C. The gut-brain axis: interactions between enteric microbiota, central and enteric nervous systems. Ann Gastroenterol. 2015;28:203–9. https://pubmed.ncbi.nlm.nih.gov/25830558.

7. Sekirov I, Russell SL, Antunes LCM, Finlay BB. Gut microbiota in health and disease. Physiol Rev. 2010;90:859–904. doi:10.1152/physrev.00045.2009.

8. Boulangé CL, Neves AL, Chilloux J, Nicholson JK, Dumas M-E. Impact of the gut microbiota on inflammation, obesity, and metabolic disease. Genome Med. 2016;8:42. doi:10.1186/s13073-016-0303-2.

9. Kamada N, Seo S-U, Chen GY, Núñez G. Role of the gut microbiota in immunity and inflammatory disease. Nat Rev Immunol. 2013;13:321–35. doi:10.1038/nri3430.

10. Mescher MF, Strominger JL, Watson SW. Protein and carbohydrate composition of the cell envelope of Halobacterium salinarium. J Bacteriol. 1974;120:945–54.

11. Silhavy TJ, Kahne D, Walker S. The bacterial cell envelope. Cold Spring Harb Perspect Biol. 2010;2:a000414–a000414. doi:10.1101/cshperspect.a000414.

12. Beveridge TJ. Structures of gram-negative cell walls and their derived membrane vesicles. J Bacteriol. 1999;181:4725–33. doi:10.1128/JB.181.16.4725-4733.1999.

13. Whitfield C. Bacterial extracellular polysaccharides. Can J Microbiol. 1988;34:415–20. doi:10.1139/m88-073.

14. Emi Y, Hiroaki T, Jun H, Tohru I, Tomoyuki S. Lectin microarray reveals binding profiles of Lactobacillus casei strains in a comprehensive analysis of bacterial cell wall polysaccharides. Appl Environ Microbiol. 2011;77:4539–46. doi:10.1128/AEM.00240-11.

15. Minoshima F, Ozaki H, Odaka H, Tateno H. Integrated analysis of glycan and RNA in single cells. bioRxiv. 2021;:2020.06.15.153536. doi:10.1101/2020.06.15.153536.

16. Hevia A, Delgado S, Margolles A, Sánchez B. Application of density gradient for the isolation of the fecal microbial stool component and the potential use thereof. Sci Rep. 2015;5:16807. doi:10.1038/srep16807.

17. Klindworth A, Pruesse E, Schweer T, Peplies J, Quast C, Horn M, et al. Evaluation of general 16S ribosomal RNA gene PCR primers for classical and next-generation sequencing-based diversity studies. Nucleic Acids Res. 2013;41:e1–e1. doi:10.1093/nar/gks808.

18. Bolyen E, Rideout JR, Dillon MR, Bokulich NA, Abnet CC, Al-Ghalith GA, et al. Reproducible, interactive, scalable and extensible microbiome data science using QIIME 2. Nat Biotechnol. 2019;37:852–7. doi:10.1038/s41587-019-0209-9.

19. Callahan BJ, McMurdie PJ, Rosen MJ, Han AW, Johnson AJA, Holmes SP. DADA2: High-resolution sample inference from Illumina amplicon data. Nat Methods. 2016;13:581–3. doi:10.1038/nmeth.3869.

20. Bokulich NA, Kaehler BD, Rideout JR, Dillon M, Bolyen E, Knight R, et al. Optimizing taxonomic classification of marker-gene amplicon sequences with QIIME 2’s q2-feature-classifier plugin. Microbiome. 2018;6:90. doi:10.1186/s40168-018-0470-z.

21. Quast C, Pruesse E, Yilmaz P, Gerken J, Schweer T, Yarza P, et al. The SILVA ribosomal RNA gene database project: Improved data processing and web-based tools. Nucleic Acids Res. 2013;41 Database issue:D590–6. doi:10.1093/nar/gks1219.

22. Robeson M, O’Rourke D, Kaehler B, Ziemski M, Dillon M, Foster J, et al. RESCRIPt: Reproducible sequence taxonomy reference database management for the masses. 2020. doi:10.1101/2020.10.05.326504.

23. Catherine L, Rob K. UniFrac: a New Phylogenetic Method for Comparing Microbial Communities. Appl Environ Microbiol. 2005;71:8228–35. doi:10.1128/AEM.71.12.8228-8235.2005.

24. Claesson MJ, Cusack S, O’Sullivan O, Greene-Diniz R, de Weerd H, Flannery E, et al. Composition, variability, and temporal stability of the intestinal microbiota of the elderly. Proc Natl Acad Sci U S A. 2011;108 Suppl Suppl 1:4586–91. doi:10.1073/pnas.1000097107.

25. Low A, Soh M, Miyake S, Seedorf H. Host-age prediction from fecal microbiome composition in laboratory mice. bioRxiv. 2020;:2020.12.04.412734. doi:10.1101/2020.12.04.412734.

26. Varki A. Sialic acids in human health and disease. Trends Mol Med. 2008;14:351–60. doi:10.1016/j.molmed.2008.06.002.

27. Almagro-Moreno S, Boyd EF. Insights into the evolution of sialic acid catabolism among bacteria. BMC Evol Biol. 2009;9:118. doi:10.1186/1471-2148-9-118.

28. Bäckhed F, Roswall J, Peng Y, Feng Q, Jia H, Kovatcheva-Datchary P, et al. Dynamics and stabilization of the human gut microbiome during the first year of life. Cell Host Microbe. 2015;17:852. doi:https://doi.org/10.1016/j.chom.2015.05.012.

29. Zhang C, Li S, Yang L, Huang P, Li W, Wang S, et al. Structural modulation of gut microbiota in life-long calorie-restricted mice. Nat Commun. 2013;4:2163. doi:10.1038/ncomms3163.

30. Pan F, Zhang L, Li M, Hu Y, Zeng B, Yuan H, et al. Predominant gut Lactobacillus murinus strain mediates anti-inflammaging effects in calorie-restricted mice. Microbiome. 2018;6:54. doi:10.1186/s40168-018-0440-5.

31. Yatsunenko T, Rey FE, Manary MJ, Trehan I, Dominguez-Bello MG, Contreras M, et al. Human gut microbiome viewed across age and geography. Nature. 2012;486:222–7. doi:10.1038/nature11053.

32. Claesson MJ, Jeffery IB, Conde S, Power SE, O’Connor EM, Cusack S, et al. Gut microbiota composition correlates with diet and health in the elderly. Nature. 2012;488:178–84. doi:10.1038/nature11319.

33. van Tongeren SP, Slaets JPJ, Harmsen HJM, Welling GW. Fecal microbiota composition and frailty. Appl Environ Microbiol. 2005;71:6438–42. doi:10.1128/AEM.71.10.6438-6442.2005.

34. Sakellaris G, Kolisis FN, Evangelopoulos AE. Presence of sialic acids in Lactobacillus plantarum. Biochem Biophys Res Commun. 1988;155:1126–32. doi:https://doi.org/10.1016/S0006-291X(88)81257-9.

35. Varki A, Gagneux P. Multifarious roles of sialic acids in immunity. Ann N Y Acad Sci. 2012;1253:16–36. doi:10.1111/j.1749-6632.2012.06517.x.

36. Chang Y-C, Nizet V. Siglecs at the host-pathogen interface. Adv Exp Med Biol. 2020;1204:197–214. doi:10.1007/978-981-15-1580-4_8.

37. Angata T, Varki A. Chemical diversity in the sialic acids and related α-keto acids: An evolutionary perspective. Chem Rev. 2002;102:439–70. doi:10.1021/cr000407m.

